# C_4_ photosynthesis and climate through the lens of optimality

**DOI:** 10.1101/048900

**Authors:** Haoran Zhou, Brent R. Helliker, Erol Akçay

**Affiliations:** Department of Biology, University of Pennsylvania, Philadelphia, PA 19104, USA

**Keywords:** C_4_ evolution, optimal stomatal conductance, resource allocation, water limitation, selective pressure, dark/light reaction

## Abstract

CO_2_, temperature, water availability and light intensity were all potential selective pressures to propel the initial evolution and global expansion of C_4_ photosynthesis over the last 30 million years. To tease apart how the primary selective pressures varied along this evolutionary trajectory, we coupled photosynthesis and hydraulics models while optimizing photosynthesis over stomatal resistance and leaf/fine-root allocation. We further examined the importance of resource (e.g. nitrogen) reallocation from the dark to the light reactions during and after the initial formation of C_4_ syndrome. We show here that the primary selective pressures—all acting upon photorespiration in C_3_ progenitors—changed through the course of C_4_ evolution. The higher stomatal resistance and leaf-to-root allocation ratio enabled by the C_4_ carbon-concentrating mechanism led to a C_4_ advantage without any change in hydraulic properties, but selection on nitrogen reallocation varied. Water limitation was the primary driver for the initial evolution of C_4_ 25-32 million years ago, and could positively select for C_4_ evolution with atmospheric CO_2_ as high as 600 ppm. Under these high CO_2_ conditions, nitrogen reallocation was necessary. Low CO_2_ and light intensity, but not nitrogen reallocation, were the primary drivers during the global radiation of C_4_ 5-10 MYA. Finally, our results suggest that identifying the predominate selective pressures at the time C_4_ first evolved within a lineage should help explain current biogeographical distributions.

**Statement of authorship:** HZ, BH and EA conceptualized the study. HZ and EA built the model, HZ and BH put the idea in a general evolutionary context, HZ performed the modeling work and analyzed output data. HZ wrote the first draft, BH and EA contributed substantially to revisions.

**Significance Statement:** C_4_ photosynthesis pathway had evolved more than 60 times independently across the terrestrial plants through mid-Oligocene (~30 MYA) and diversified at late Miocene (5 to 10 MYA). We use an optimal physiology model to examine the primary selective pressures along the evolutionary history. Water limitation was the primary driver for C_4_ evolution from the initial evolutionary events 25-32 MYA until CO_2_ became low enough to, along with light intensity, drive the global radiation of C_4_ 5-10 MYA. This modeling framework can be used to investigate evolution of other physiological traits (e.g. N reallocation, hydraulics) after the initial formation of C_4_ syndrome, which contributed to further increasing productivity of C_4_ in historical and current environmental conditions.

## Introduction

The evolution of the C_4_ photosynthetic pathway enabled the concentration of CO_2_ around Rubisco, the enzyme responsible for the first major step of carbon fixation in the C_3_ photosynthetic pathway, thus eliminating photorespiration. C_3_ photosynthesis is present in all plants, and within C_4_ plants the C_3_ pathway is typically ensconced within specialized bundle sheath cells that surround leaf veins. CO_2_ that diffuses into a leaf is shuttled from adjacent mesophyll cells to the bundle sheath via a four-carbon pump, the energetic cost of which is remunerated by ATP derived from the light reactions (1, 2). As a whole, the C_4_ pathway eliminates photorespiration, a process that can dramatically reduce photosynthesis and begins with the assimilation of O_2_, instead of CO_2_, by Rubisco. Over the last 30 million years, the reduction in C_3_ photosynthesis by photorespiration was large and broad enough to select for the independent evolution of the C_4_ pathway more than 60 times across the terrestrial plants (3). The diversity of plant families with C_4_ is greatest in the eudicots (1200 species) and the Poaceae, the monocot family containing the grasses (4500 species) (2), account for nearly 25% of terrestrial plant productivity and several important agricultural species (4).

While increased photorespiration in C_3_ progenitors was central to the evolution of the C_4_ carbon concentrating mechanism (CCM), the relative importance of different environmental drivers of the photorespiratory increase has been the subject of much debate (5, 6, 7, 8). Lower CO_2_ leads to higher rates of photorespiration, as does higher temperature. Past physiological models therefore focused on examining temperature and CO_2_ concentration as selective pressures for C_4_ evolution and expansion (5, 9, 10). Under warmer temperatures and low CO_2_, C_4_ photosynthesis has greater carbon gain than C_3_, but under cooler temperatures and high CO_2_, the metabolic costs of the C_4_ pathway and lower photorespiration in C_3_ leads to greater carbon gain in C_3_. Alternatively, water availability has been proposed as the impetus for C_4_ evolution in eudicots (2), and recent phylogenetic analyses have suggested the same in grasses (7, 13). Water availability should have an impact on C_4_ evolution that could work independently or in concert with changes in CO_2_ and temperature. First, water deficits indirectly increase photorespiration in C_3_ plants by forcing stomatal closure to reduce leaf water loss; consequently decreasing the flux of CO_2_ into the leaf and the availability of CO_2_ for Rubisco. Second, the C_4_ carbon concentrating mechanism allows for the maintenance of lower stomatal conductance, and therefore lower water loss, for a given assimilation rate; leading to a higher water-use-efficiency (WUE) than C_3_ (11, 12).

The different environmental drivers of the photorespiratory increase in C_3_ plants—atmospheric CO_2_ concentration, temperature and water availability—have changed dramatically as C_4_ photosynthesis has evolved over the last 30 million years. Atmospheric CO_2_ decreased monotonically from the mid-Oligocene (680 ± 200 ppm) to the early Miocene (357 ± 108 ppm) down to the Pleistocene minima, where CO_2_ oscillated between approximately 180 and 280 ppm through glacial/interglacial cycles (14). Physiological models that focused on temperature and CO_2_ implied that C_4_ evolved, in both grasses and eudicots, at the low end of this CO_2_ range in the mid-Miocene to the Pleistocene (2, 5, 9, 10, 15). C_4_ grasses did become a major component of grassland biomes—in terms of biomass, C_4_ lineage diversity, or herbivore dietary components—in the late Miocene (5 to 10 MYA), but molecular evidence suggests that C_4_ photosynthesis arose in the grasses in the mid-Oligocene (~30 MYA) (16). Similarly, phylogenetic reconstructions provide evidence that eudicots of the Chenopodiaceae evolved C_4_ as early as the monocots, but saw the greatest rate of C_4_ evolution and diversification in the late Miocene (17, 18, 19). Along with CO_2_, precipitation declined over the period of C_4_ evolution and diversification, leading to vast terrestrial areas where low or highly seasonal precipitation inputs led to the loss of forests and consequently, the evolution of the world’s first grasslands (20, 21). The spread of grasslands indicate a habitat change with larger surface radiation loads, higher surface temperatures and increased potential for plant water loss (6, 22). Therefore, the early evolution of C_4_ suggested by molecular phylogenies indicates that water availability played an important role for both C_4_ grasses and eudicots while CO_2_ was still relatively high (6, 18, 23, 24, 22, 25). The potentially interacting roles of water availability, changes in radiation and CO_2_ along the evolutionary trajectory of C_4_ photosynthesis have not been fully investigated within a comprehensive physiological model.

A related but largely unstudied evolutionary change during the divergence of C_4_ photosynthesis from C_3_ is the allocation of nutrients/resources (e.g. N considering enzymes and proteins) between the dark reactions and the light reactions. C_4_ plants might allocate a greater proportion of N to light reactions than to dark reactions as compared to C_3_ because of the extra ATP costs of the CCM (26, 27). We propose that the reallocation of N between dark and light reactions provides a further advantage for C_4_ above the CCM alone, and that different environmental conditions can select for a shift in the degree of reallocation both through evolutionary time and across species in extant plants.

Changes in the environmental controls on photorespiration suggest that multiple environmental drivers interacted to differing degrees along the trajectory of C_4_ evolution. Our goal here is to tease apart the selective pressures that led to the evolution of C_4_ photosynthesis initially and the global expansion 5-10 MYA through to the current day. We use the framework of an optimality model in which the plant makes allocation “decisions” in order to maximize photosynthetic assimilation rate. Our approach advances our understanding of C_4_ evolution in four important ways. First, we revisit the temperature-CO_2_ crossover approach and integrate the effects of water limitation, light, optimal allocation decisions, and the interactions between these in a single model. Second, the hypothesis that C_4_ photosynthesis has a higher WUE than C_3_ implicitly relies on an optimality argument to balance carbon gain and water loss (28), yet the role of optimal stomatal conductance in mediating selective pressures due to water limitation during the evolution of C_4_ plants remains largely unexplored (but see 15). Most previous models assume *a priori* that C_4_ grasses have lower stomatal conductance. Instead, we let both stomatal conductance and leaf/fine-root allocation emerge endogenously from the model. Third, we use the C_4_ model including cost of the C_4_ pathway in the light reactions (2 additional ATP per CO_2_ fixed; 1, 29), which previous modeling analysis did not explicitly consider (9, 22, 30, 31).

Finally, we consider reallocation of nitrogen from the dark reactions to the light reactions, which can change the tradeoffs between photosynthesis and water use by C_4_ grasses.

## Results

Assimilation-based crossover temperatures, defined as the temperature at which assimilation by the C_4_ pathway exceeds that of the C_3_ pathway, decrease as water limitation increases and light intensity increases across all CO_2_ concentrations (Fig. 1, Fig. S1). Without water stress (solid black line in Fig 1), our model predicts a C_3_/C_4_ crossover temperature of 23°C under 380 ppm; a result similar to previous data and/or models that did not explicitly account for water stress (9, 10, 32). The model results in Fig. 1 were all under saturated light and with a C_4_ *J*_*max*_/*V*_*cmax*_ ratio of 4.5, which corresponds to a full reallocation of nitrogen from light to dark reactions. Model results for a C_4_ *J*_*max*_/*V*_*cmax*_ ratio of 2.1 (corresponding to no reallocation) were similar (Fig. S1a) with the primary exception being that under low CO_2_ and low water availability (e.g. CO_2_=300 ppm, VPD = 3 kPa and Ψ_S_ = -1.5 MPa or all CO_2_ concentrations with higher VPD and lower Ψ_S_, crossover temperatures are higher with *J*_*max*_/*V*_*cmax*_ = 4.5, showing that nitrogen reallocation decreases the C_4_ advantage under water limitation and low CO_2_. Under saturated soil and low VPD, crossover temperatures decrease along with increasing light intensity (Fig. S1c, d). An increase in light intensity provides a larger relative benefit for C_4_ at low CO_2_, because C_3_ photosynthesis remains CO_2_ limited throughout while C_4_ light limitations lessen as light increases.

**Fig. 1.**
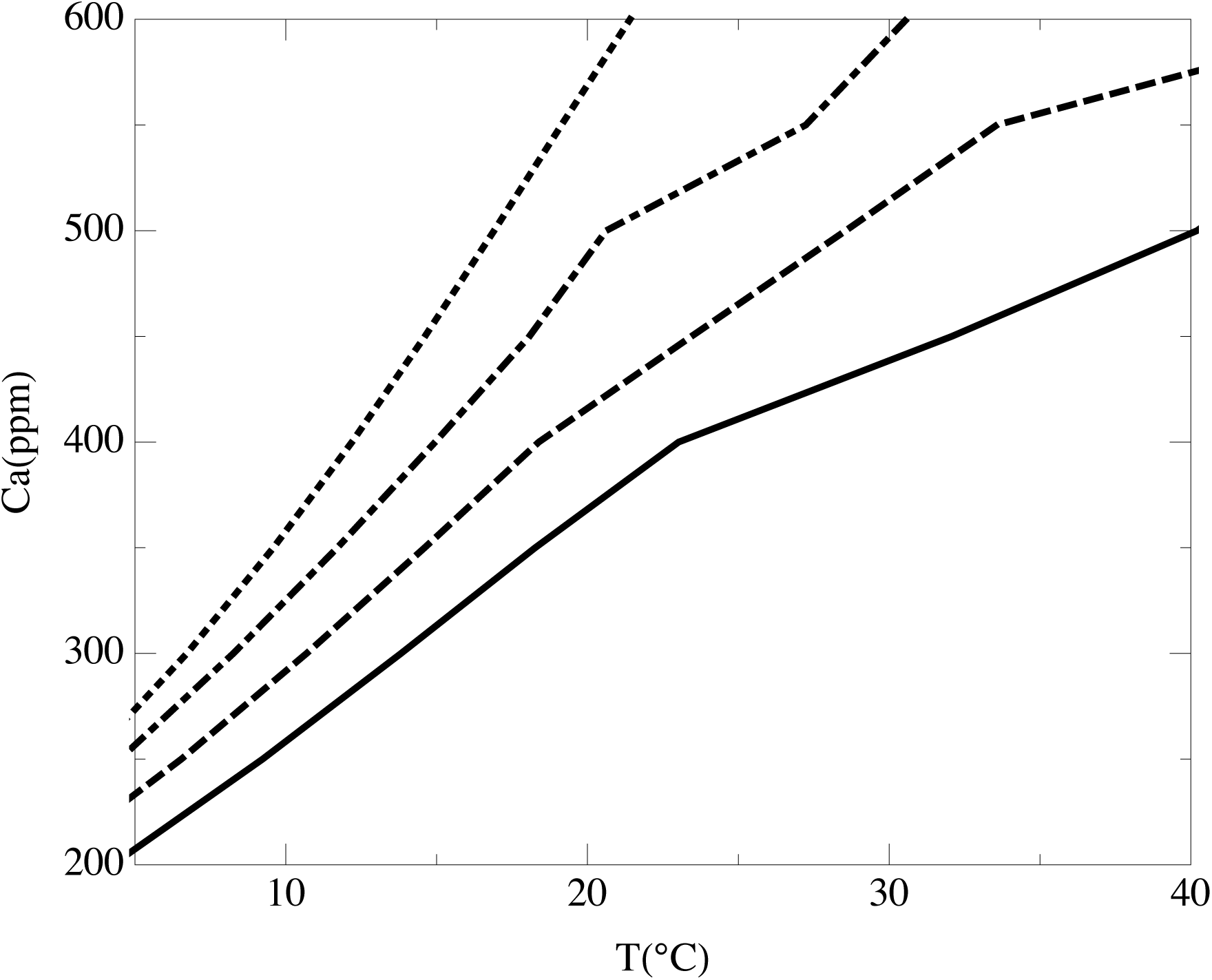
Crossover temperatures of photosynthesis for C_3_ and C_4_ with the change of CO_2_ concentration under different water conditions. Light intensity was 1400 μmol photons m^-2^s^-1^ for all model runs. *J*_max_/*V*_cmax_=2.1 for C_3_ and *J*_max_/*V*_cmax_=4.5 for C_4_. Solid black line: VPD=0.15kPa, Ψ_S_=0 MPa; dashed black line: VPD=1kPa, Ψ_S_=-0.5 MPa; dot-dashed black line: VPD=2kPa, Ψ_S_=-1 MPa; dotted black line: VPD=3 kPa, Ψ_S_=-1.5 MPa.

While crossover temperatures allow for a clear diagnostic of comparative assimilation, they do not demonstrate the degree of C_4_ photosynthetic advantage. To this end, we calculated the net assimilation rate difference between C_4_ and C_3_, Δ*A*_*n*_ (net assimilation of C_4_ minus that of C_3_), to comprehensively examine the whole suite of environmental conditions (Fig. 2, 3). The positive contour space (Δ*A*_*n*_ > 0) means that C_4_ outcompetes C_3_ within given environmental dimensions, and the higher the Δ*A*_*n*_, the greater the advantage of C_4_. Under a CO_2_ concentration of 200 ppm and saturated light, Δ*A*_*n*_ is higher under moist conditions than water-limited conditions (Fig. 2a, b). In contrast, under higher CO_2_ concentrations (400 and 600 ppm), C_4_ has the greatest advantage only in water-limited conditions, which leaves a relatively small environmental envelope for C_4_ to evolve (areas where Δ*A*_*n*_ >0 in Fig. 2c-f). This result is due to the fact that C_3_ photosynthesis has a greater proportional increase in assimilation from 200 to 400 and 600 ppm CO_2_. Across all scenarios, increasing *J*_*max*_/*V*_*cmax*_ increases both the Δ*A*_*n*_ and space for C_4_ advantage (Fig. 2 b, d, f). At 200 ppm and saturated soils, Δ*A*_*n*_ is highest under saturated light, and decreases as light intensity decreases (Fig. 3a, b). At 400 ppm CO_2_ and higher, Δ*A*_*n*_ indicated a C_4_ advantage (Δ*A*_*n*_ > 0) only when light intensities were above 1000 mmol m-2 s-1, temperature was above 30 °C and *J*_*max*_/*V*_*cmax*_ was high (Fig. 3 c, d). Finally, we calculated the photosynthesis rates of the two pathways under conditions often encountered in today’s grasslands to look at the effect of nitrogen reallocation (Fig. 4). With *J*_*max*_/*V*_*cmax*_ =2.1 for both C_3_ (solid black line) and C_4_ (dashed line), the C_4_ assimilation rate is rarely higher than C_3_, which indicates C_4_ does not have an obvious advantage under current CO_2_ from saturated soils down to VPD = 2 kPa and Ψ_S_ = -1 MPa. However, with *J*_*max*_/*V*_*cmax*_ =4.5 for C_4_ (dotted line), C_4_ does have an advantage over C_3_ at temperatures above 25°.

**Fig. 2.**
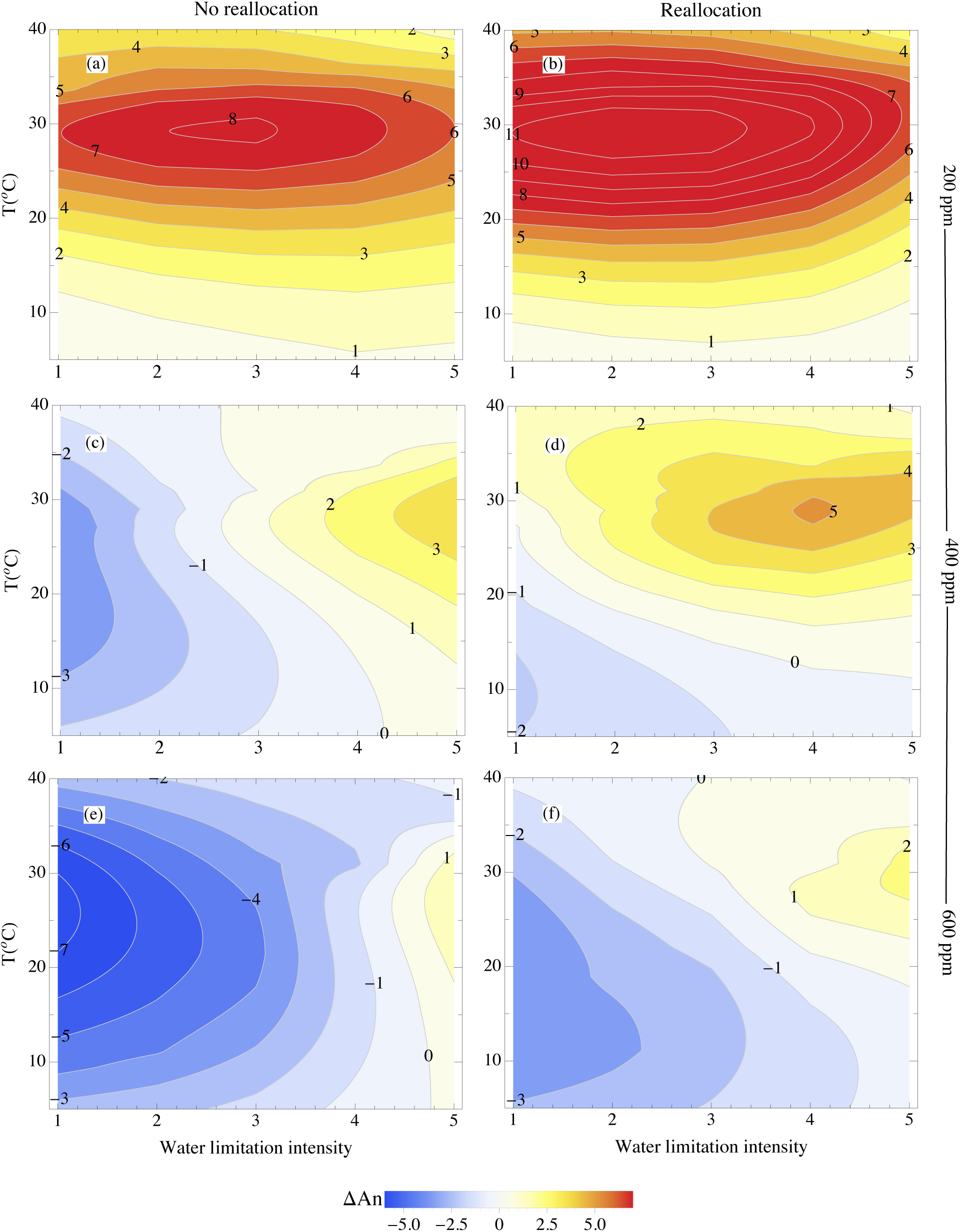
The total difference in CO_2_ assimilation between C_4_ and C_3_ (*A*_n_(C_4_)-*A*_n_(C_3_)) under various CO_2_ (200 ppm, 400 ppm and 600 ppm) and water conditions under saturated light intensity (1400 μmol photons m^-2^s^-1^). *J*_max_/*V*_cmax_=2.1 for C_3_ and C_4_ (a, c, e) and *J*_max_/*V*_cmax_=2.1 for C_3_ and *J*_max_/*V*_cmax_=4.5 for C_4_ (b, d, f). Water limitation intensity is: 1: VPD =0.15 kPa, Ψ_S_=0 MPa; 2: 1.5 kPa, -0.5MPa; 3: 2kPa, -1 MPa; 4: 3kPa, -1.5 MPa; 5: 4kPa, -2 MPa.

**Fig. 3.**
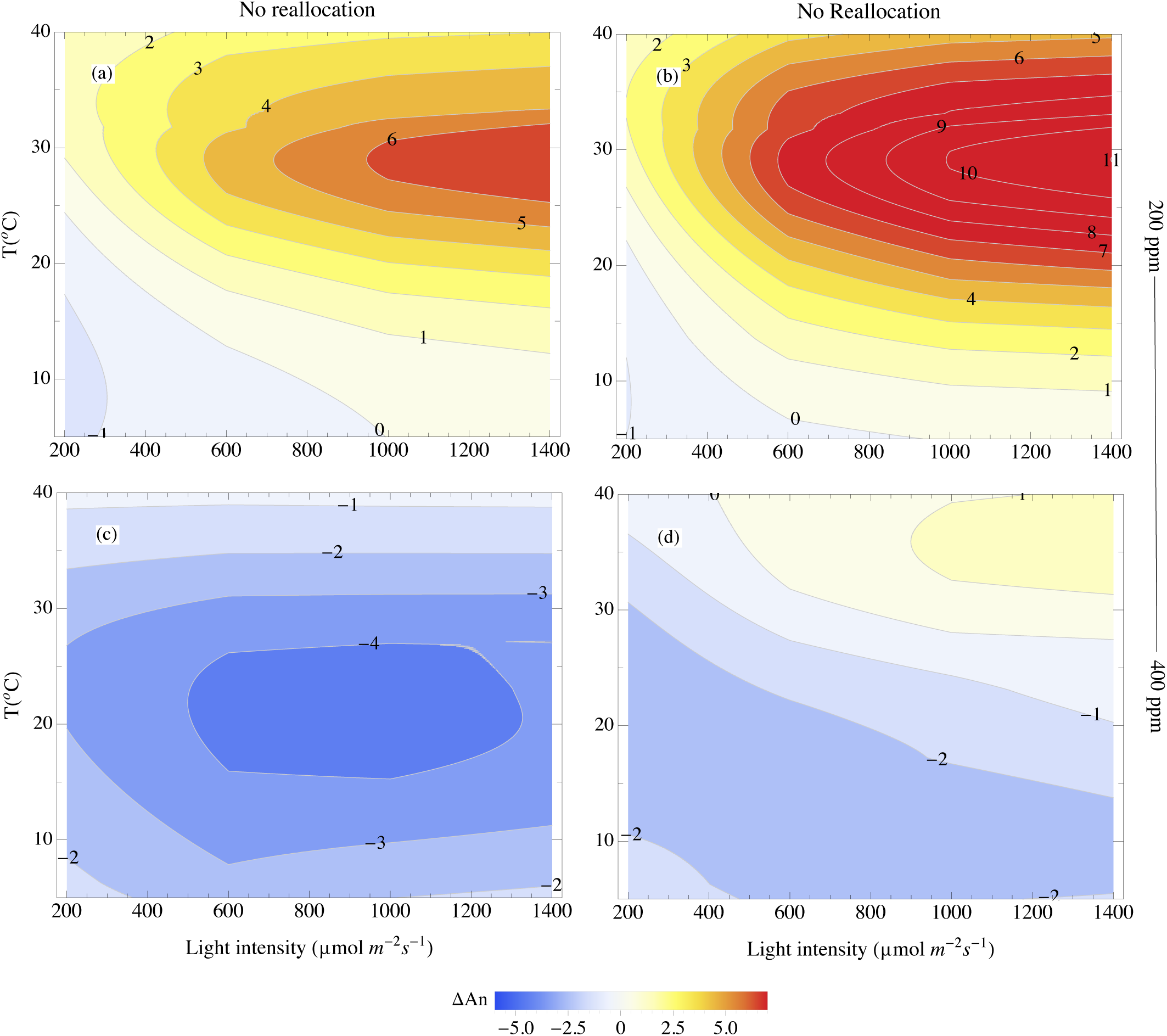
The total difference in CO_2_ assimilation between C_4_ and C_3_ (*A*_n_(C_4_)-*A*_n_(C_3_)) with *J*_max_/*V*_cmax_=2.1 for C_3_ and C_4_ under various CO_2_ (200 ppm, 400 ppm) and different light intensities (from 200 to 1400 μmol photons m^-2^s^-1^) with saturated water condition (VPD=0.15kPa, Ψ_S_=0 MPa) (a, c) and with *J*_max_/*V*_cmax_=2.1 for C_3_ and *J*_max_/*V*_cmax_=4.5 for C_4_ (b, d).

**Fig. 4.**
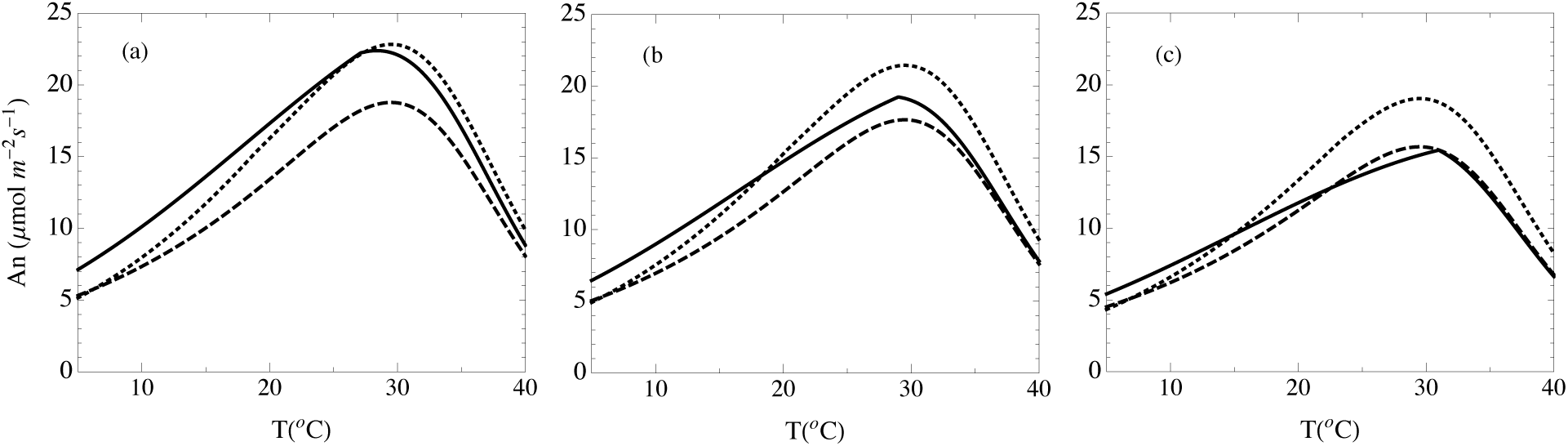
Assimilation rates of C_3_ with *J*_max_/*V*_cmax_=2.1 (solid black line), C_4_ with *J*_max_/*V*_cmax_=2.1 (dashed black line) and C_4_ with *J*_max_/*V*_cmax_=4.5 (dotted black line) under light intensity of 1400 μmol photons m^-2^s^-1^, CO_2_ of 400 ppm and different water limitation conditions. (a) saturated soils; (b) VPD=1kPa and Ψ_S_-0.5 MPa; (c) VPD=2kPa and Ψ_S_=-1 MPa.

Under all environmental and nitrogen allocation scenarios, optimal stomatal resistance (*r*_*s*_) and leaf biomass/total biomass allocation (f) are higher in C_4_ plants than C_3_ plants. The general pattern of response was similar across CO_2_ concentrations, so only 400 ppm is presented with both highest and lowest water availability (Fig. 5 and Fig. S2). Optimal *f* and *r*_*s*_ for C_3_ is always lower than that for C_4_ under different water availability and CO_2_ (Fig. 5). In addition, *f* decreases and *r*_*s*_ *increases* as intensity of water limitation increases. Results are the consistent for C_4_ with a *J*_*max*_/*V*_*cmax*_ of 2.1 and *J*_*max*_/*V*_*cmax*_ of 4.5. The jumps in r_s_ and *f* in Fig. 5 correspond to the transition from RuBP carboxylation limited assimilation to RuBP regeneration limited assimilation (*A*_*j*_). The transition temperature decreases as CO_2_ increases. In the Fig. S3, the various limitation states are plotted together under multiple environmental scenarios, using both *J*_*max*_/*V*_*cmax*_*=2.1* and 4.5 for C_4_. With *J*_*max*_/*V*_*cmax*_*=2.1*, C_4_ is light limited in most of the environmental conditions. With *J*_*max*_/*V*_*cmax*_=4.5, C_4_ starts to be limited by CO_2_ under low temperatures and to be limited by light under high temperatures.

**Fig. 5.**
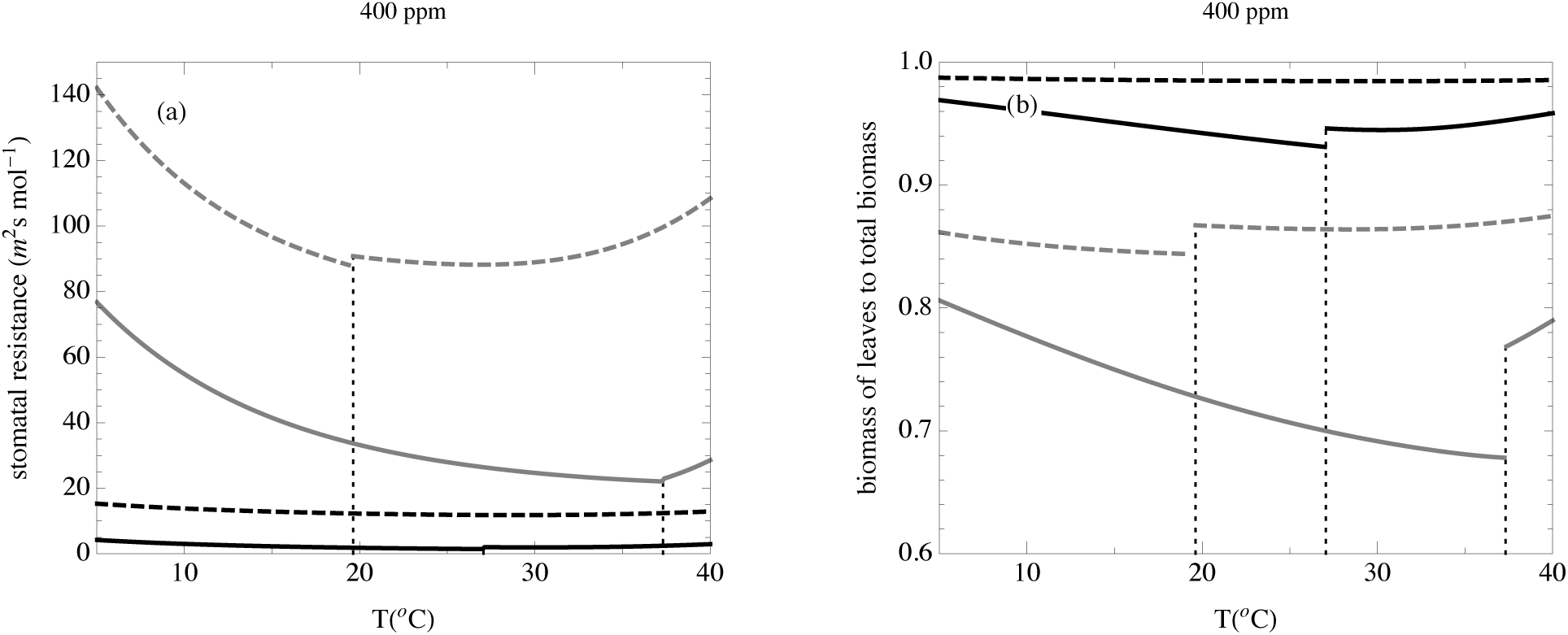
Stomatal resistance (r_s_) and leaf/fine-root allocation (f) as a function of temperature, with *J*_max_/*V*_cmax_=2.1 for both C_3_ and C_4_ with saturated light under CO_2_ of 400 ppm and different water conditions. Solid black line: C_3_ with VPD=0.15kPa, Ψ_S_=0 MPa; dashed black line: C_4_ with VPD=0.15kPa, Ψ_S_=0; solid grey line: C_3_ with VPD=4 kPa, Ψ_S_=-2 MPa; dashed grey line: C_4_ with VPD=4 kPa, Ψ_S_=-2 MPa. Vertical lines indicate transition from RuBP carboxylation limited condition to RuBP regeneration limited condition for C_3_ and C_4_.

## Discussion

Our modeling results imply that the environmental selection for C_4_ evolution was in place during the mid-Oligocene at warm, arid sites as water limitation acted as the primary selective pressure to increase photorespiration when CO_2_ is above 400 ppm, and even up to 600 ppm. Under saturated water conditions, there was little room for C_4_ to evolve 20-30 MYA as CO_2_ was likely above 400 ppm (14, 33), and the ATP costs of the C_4_ mechanism is too high and photorespiration in C_3_ plants too low. As water becomes more limited, however, the C_4_ advantage becomes increasingly larger. Enhanced carbon gain under water limited conditions has been believed to be the selective force behind the evolution of C_4_ in dicotyledonous plants (2) and more recently, in accordance with phylogenetic evidence, in grasses (7, 16). Molecular-based evidence supports a mid-Oligocene evolution of C_4_ in some grass and eudicot lineages, and our results suggest the same (8, 18, 34). Lineages that evolved in C_4_ during this period may have been predisposed to exist as arid-land, saline, or disturbed-site specialists as we see today in most C_4_ eudicots and the earliest known grass subfamily in which C_4_ evolved, the Chloridoideae (8, 35).

As CO_2_ decreased through the Miocene, warm temperatures remained a strong selective force, but the main selective force for C_4_ evolution shifted from water limitation to low CO_2_ and, to a lesser extent, light intensity. Since increased light intensity alone could not lead to advantage of C_4_ under high CO_2_ (Fig. 3c), it seems likely that C_4_ grasses could not dominate open grasslands, except in locally arid areas, while CO_2_ was still high. So between the initial evolutionary events leading to the emergence of C_4_ and the large-scale expansion within the grasses 5-10 MYA, C_4_ grass radiation likely idled in small pockets of selective favorability as CO_2_ concentrations declined through the Miocene (2, 8). As CO_2_ declined, the high light levels inherent to grassland systems gave C_4_ photosynthesis an increasing advantage, leading to broader geographic and evolutionary radiation. Our results are consistent with previous studies showing that low CO_2_ (200-300 ppm) strongly favors C_4_ over C_3_ photosynthesis (e.g. 9, 15). And we further show that low CO_2_ provides a clear C_4_ advantage under a large range of water availability and light intensity regimes. The greatest C_4_ advantage occurs, however, in relatively moist and mildly water-limited conditions; opposite to that which is seen under high CO_2_ (Fig. 2c). The environmental conditions that lead to the largest C_4_ advantage within our model therefore perfectly parallels those documented in the 20^th^ century, C_4_-dominated grasslands: highly seasonal precipitation that occurs chiefly within a warm growing season (36, 37, 38). These are also similar to the conditions that led to the large-scale expansion of C_4_ grasslands in the Miocene, for example the onset of summer monsoons and subsequent C_4_ grassland expansion in the Indian subcontinent (39).

The potential role of water limitation to play a central role C_4_ grass evolution has sparked interest in grass hydraulics and the anatomical shifts in C_3_ grasses that were prerequisites to C_4_ evolution (24, 22, 40). The modeling effort of Osborne and Sack (22) suggests a hydrological underpinning to the evolution of C_4_ grasses, but found a much smaller environmental window for C_4_ evolution than we did. At 400 ppm CO_2_ and soil water potential of -1 MPa—a common occurrence in grasslands (41)—they showed that C_4_ hydraulic conductance must be twice that of C_3_ grasses for C_4_ grasses to achieve slightly greater carbon uptake. In contrast, we find a clear C_4_ advantage under these, and even drier, conditions by allowing for optimal solutions of stomatal conductance and leaf/fine-root ratio to maximize photosynthesis. Plant hydraulic conductance was kept equal across C_3_ and C_4_ throughout simulations, and increasing hydraulic conductance had little impact on our major results and conclusion (Fig. S4, S5), the implication being that the C_4_ pathway itself is enough to result in greater carbon gain under water stress without any required increase in hydraulic conductance. These results do not necessarily contradict the idea that larger bundle sheaths and smaller interveinal distance—which were clear prerequisites for C_4_ evolution (24, 42) — led to greater hydraulic conductance and drought tolerance among C_3_ grass progenitors (24), but they do suggest that greater hydraulic conductance is not necessary to give C_4_ plants an advantage once the carbon-concentrating mechanism evolved. In fact, we hypothesize that once C_4_ evolves in a lineage, selection on increased hydraulic conductance would not only lessen, but invert, leading to the development of even narrower xylem conduits and greater drought resistance. There is some empirical support for such a prediction in eudicots (43).

We assumed that during the early evolution of C_4_, both C_3_ and C_4_ plants had a similar balance of nitrogen across the light and dark reactions, and that the allocation of nitrogen could be treated separately from the evolution of the C_4_ CCM as a target of selection. We propose that different environmental conditions can select for a shift in the degree of reallocation (assessed here by a change in *J*_*max*_/*V*_*cmax*_) both through evolutionary time and across species in extant plants. In general, CCMs allow for less investment in nitrogen-rich Rubisco (44), and the nitrogen not used for Rubisco could be either reinvested in light harvesting machinery, or simply not used at all; thus reducing the plant nitrogen requirement. Modeling studies have long assumed a high *J*_*max*_/*V*_*cmax*_ for C_4_ photosynthesis (22, 31, 45) and measurements show lower Rubisco content and higher chlorophyll and thylakoid content, giving evidence of reallocation in extant C_4_ species (26, 27, 46). Empirical estimates of *J*_*max*_/*V*_*cmax*_, in C_4_ plants paint a more variable picture, ranging from 2 to above 6, with a mean of around 4.5 (47, 48, 49, 50, 51, 52), which is higher than the mean *J*_*max*_/*V*_*cmax*_ estimates for C_3_ plants of 2.1 (53). Increasing *J*_*max*_/*V*_*cmax*_ almost always increases the photosynthesis rate of C_4_ grasses (Fig. 4, Fig. S6), and therefore could lead to a competitive advantage over C_3_ grasses as well as C_4_ grasses that do not reallocate. Assuming there is little cost or no genetic constraints for reallocation, the selection pressure to reallocate would have been strongest when CO_2_ was high, e.g. during the initial evolutionary events in the Oligocene/Miocene, when the CCM alone does not give C_4_ a large advantage (Fig. 2c, e, Fig. 3c). When CO_2_ was low during the C_4_ radiation 5-10 MYA, however, the CCM alone would give C_4_ an advantage and reallocation would not change the competitive balance between C_3_ and C_4_ (Fig. 2a and Fig. 3a). As CO_2_ remained low through to the Pleistocene, selection for nitrogen reallocation to the light reactions would lessen further, especially during the CO_2_ minima of the Pleistocene glacial periods (~180 ppm). In this context, an interesting question is how the evolutionary picture of *J*_*max*_ and *V*_*cmax*_ allocation was coordinated with the formation of C_4_ pathway in response to the high CO2 in Oligocene, to the CO2 decrease through Pleistocene and, further, to the increase of CO_2_ in the last 150 years.

C_4_ photosynthesis first evolved 25 – 32 MYA, and many subsequent and independent evolutionary origins occurred through to the Pleistocene 2.8 MYA. Each evolutionary origin represents both different selective pressures and taxonomic (genetic) constraints as climate and CO_2_ changed. Taking the Chloridoideae as an example, we can use our model to develop hypotheses along the evolutionary trajectory of C_4_ in this grass subfamily. The initial evolution of C_4_ photosynthesis 25 – 32 MYA while CO_2_ was high was driven by aridity, acting to decrease stomatal conductance that increased photorespiration in C_3_ progenitors initially, and led to higher water use efficiency upon the evolution of the CCM. There would have been enough of a reduction in water use that selection for increased hydraulic conductance would relax, allowing for the development of more resilient—and less conductive—xylem. Also at this point, there would have been strong selection for reallocation of nitrogen from the dark reactions to the light reactions. The large radiation of C_4_ within the Chloridoideae that occurred 5 – 10 MYA was likely driven by low CO_2_ and high light, and the previously-evolved hydraulic resilience would lead to this subfamily becoming dry-site specialists observed in current-day distributions (35). There would have been much less selective pressure to reallocate N during the large radiation, but such a reorganization was likely already in place within the clade. In contrast, for the lineages that first evolved C_4_ in the late Miocene (e.g. *Stipagrostis*, *Eriachne*, *Neurachne*), CO_2_ would have been the primary impetus for C_4_ evolution, but for these lineages there would have been little selection to reallocate nitrogen until the dawn of the industrial revolution. We would also expect these more recently evolved lineages to have greater hydraulic conductance than those of the Chloridoideae. By optimizing carbon gain over water loss, we developed a plausible physiological explanation for the early evolution of C_4_ and further proposed hypotheses about how the variety of traits that comprise the C_4_ syndrome developed in concert with the climate changes that occurred through the evolutionary trajectory (54). By selecting extant species within select lineages, these hypotheses can be examined empirically, ultimately providing an integrative view of the selection pressures that led to the current physiologies and distribution of C_4_ plants.

## Materials and Methods

### Overview of the model

We used different modeling scenarios to examine the advantage of C_4_ photosynthesis for the initial origin, expansion and current distribution. Initially, we assume that the CCM is the only difference between C_3_ and C_4_. This comparison corresponds to two closely related species whose other traits have not had time to diverge in response to differential selection pressures. We then examine shifts in N allocation between the light and dark reactions of C_4_, which may have happened in subsequent divergence of C_3_ and C_4_ after the CCM evolved.

The soil-plant-air water continuum was incorporated in C_3_ photosynthesis models (55) and C_4_ models (29) to examine interactions of CO_2_, water availability, light and temperature. We used the optimality approach of Givnish (1986) (56), where C_3_ and C_4_ plants optimize stomatal resistance and leaf/fine-root allocation to balance carbon gain and water loss. A full model description is in SI in supporting information with Table S1 for parameter abbreviation and Table S2 for input parameters. The model derivation using Mathematica (Wolfram Research, Inc.) and methods for numerical solutions are in SII.

### Photosynthesis model

We are using the traditional C_3_ photosynthesis models (55) and C_4_ models (29) for the photosynthesis modeling (SI in supporting information).

### Hydraulic system

At equilibrium, the rate of water loss through transpiration equals the rate of water absorption by the roots (56):

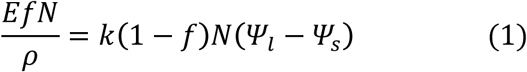

where *Ψ*_*s*_ is soil water potential, *k* is the effective root hydraulic conductivity, *N* is the total biomass of fine root and leaves, *ρ* is the leaf mass density (gcm^-2^) and *E* is the transpiration rate per leaf area. *E* could be written as *δ*/*r*_*s*_, where *δ* is the water partial pressure deficit between saturated leaf surface and the atmosphere. Thus, leaf water potential (*Ψ*_*l*_) is a function of *r*_*s*_ and leaf/fine-root allocation (*f* defined as investment into leaves/total investment in leaves and fine root)).

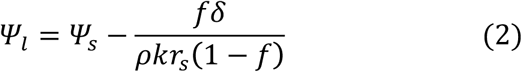

### Inhibition of photosynthesis by water stress

Reduced leaf water potential inhibits photosynthesis (57, 58, 59). We model this cost of transpiration as Weibull-type vulnerability curves relating leaf *Ψ*_*l*_ and photosynthetic parameters (45):

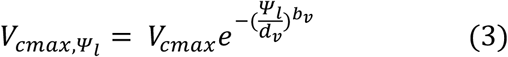

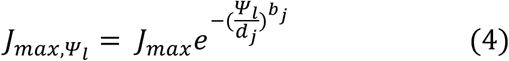

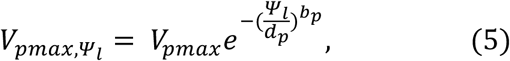

where *b* and *d* are curve fitting parameters. Since *Ψ*_*l*_ is a function of *r*_*s*_ and *f*, all those parameters are functions of *r*_*s*_ and *f*.

### Optimal stomatal resistance and optimal allocation of energy between leaves and fine roots

We assume that the plant adjusts the *r*_*s*_ and *f* to optimize the total carbon gain by

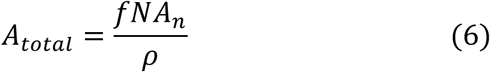

where *ρ* is the leaf mass density (g cm^-2^). As a simplifying assumption, we assume *N* and *ρ* are fixed (similar to 56). Effectively, we consider the optimization problem faced by the plant in a given instance during its growth, where its size (of which *N* is a proxy) can be regarded as a constant. Clearly, during plant growth, the assimilate will be turned into plant biomass, but the instantaneous optimization problem will still yield the optimal growth path as the growth rate is maximized at any given time. Finally, we regard *ρ* as a species-specific trait that changes at a slower time-scale than *r*_*s*_ and *f* The first order optimality conditions for *r*_*s*_ and *f* are given by (56):

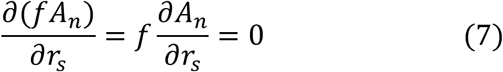

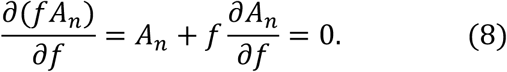

We checked the second order derivative to ensure that the numerical solutions to the first order conditions were maxima.

### Allocation of nitrogen

We examine how nitrogen allocation between RuBP carboxylation and RuBP regeneration in C_4_ grasses affect competitive advantage over C_3_ grasses. Despite great variation in *V*_*cmax*_ and *J*_*max*_ based on the total leaf nitrogen content within C_3_ plants, Wullschleger (1993) (53) found a mean of *J*_*max*_/*V*_*cmax*_ =2.1 across 109 C_3_ species, which we use as a baseline for C_3_ and C_4_ pathways in analyzing the initial evolution of C_4_. Then, we used *J*_*max*_/*V*_*cmax*_ =4.5 for C_4_ (22, 45) to analyze the role that nitrogen reallocation played in the evolutionary trajectory of C_4_ plants. In determining the values of *J*_*max*_ and *V*_*cmax*_, we used a simplified stoichiometry: we consider the total of *J*_*max*_ and *V*_*cmax*_ as a constant to hold nitrogen concentration constant (22, 45). Two assumptions are underlying this simplified stoichiometry: (1) investing one molecule of nitrogen to the dark reactions will increase of *V*_*cmax*_ equal to the increase of *J*_*max*_ by investing one molecule of nitrogen to the light reactions; (2) nitrogen allocation to enzymes involved in photorespiration and the CCM balanced each other.

### Modeling scenarios

We modeled the photosynthesis rates of C_3_ and C_4_ under temperature range from 10 °C to 40 °C with an interval of 5 °C, under CO_2_ mixing ratios ranging from 200 ppm to 600 ppm with an interval of 50 ppm, under different water conditions (VPD=0.001, 1, 2, 3, 4kPa corresponding to soil water potential (Ψ_S_ =0, -0.5, -1, -1.5, -2 MPa) and under different light intensities (1400, 1000, 600, 200, 100 μmol photons m^-2^s^-1^). We consider VPD=0.001 kPa and Ψ_s_ =0 MPa as saturated water condition and light intensity of 1400 μmol photons m^-2^ s^-1^ as an average saturated light intensity of a day.

## Acknowledgements

We are grateful for support from the University of Pennsylvania.

